# IST-editing: Infinite spatial transcriptomic editing in a generated gigapixel mouse pup

**DOI:** 10.1101/2023.12.23.573175

**Authors:** Jiqing Wu, Ingrid Berg, Viktor H. Koelzer

## Abstract

Advanced spatial transcriptomics (ST) techniques provide comprehensive insights into complex living systems across multiple scales, while simultaneously posing challenges in bioimage analysis. The spatial co-profiling of biological tissues by gigapixel whole slide images (WSI) and gene expression arrays motivates the development of innovative and efficient algorithmic approaches. Using Generative Adversarial Nets (GAN), we introduce **I**nfinite **S**patial **T**ranscriptomic **e**diting (IST-editing) and establish gene expression-guided editing in a generated gigapixel mouse pup. Trained with patch-wise high-plex gene expression (input) and matched image data (output), IST-editing enables the synthesis of arbitrarily large bioimages at inference, *e.g*., with a 106496×53248 resolution. After feeding edited gene expressions to the trained network, we model cell-, tissue- and animal-level morphological transitions in the generated mouse pup. Lastly, we discuss and evaluate editing effects on interpretable morphological features. The generated WSIs of the mouse pup and code are publicly released and accessible via https://github.com/CTPLab/IST-editing.

## Main

Recent advances in multi-omics technologies (*e.g*., spatial transcriptomics (ST)^1^) and generative artificial intelligence (AI)^2,3^ have the potential to revolutionize bioimage analysis^4^. Leveraging spatial co-profiling of high-plex mRNA transcripts (acting as proxies for gene expression) and high-resolution biomedical images, researchers possess unprecedented opportunities to model the complex spatial organization of living organisms.

Concurrently, generative AI^2,3^ has showcased remarkable progress in creating high-quality visual content, paving the way towards novel applications in the biomedical domain. Trained with Hematoxylin and Eosin (H&E)-stained or (immuno)fluorescence images, prior studies^5–7^ have achieved impressive results of bioimage generation and manipulation using GAN approaches. Recently, researchers^8^ further demonstrated the algorithmic editability on ST data and simulated cellular morphological transitions by shifting gene expression distributions. Notably, these studies were carried out at cell- or tissue-level and the generated bioimage resolution is usually smaller than 256×256. Due to scalability limitations, the generative competence of such algorithmic methods cannot be extended to the entirety of a WSI without inducing visible stitching artifacts, which can be partially mitigated by employing more hardware resources. A series of StyleGAN studies^9,10^ first showed the feasibility of training 1024×1024 images on 8 V100 GPUs. In a recent paper, a GAN-based approach^11^ has accomplished 4096×4096 image generation with remarkable fine details. Critically, this achievement was made possible by training a scaled GAN model on 96-128 A100 GPUs. Despite impressive breakthroughs in generating megapixel-resolution images, the hardware requirements for modeling WSIs at the gigapixel scale can be computationally intractable, making the application of such a model prohibitively expensive in biomedical research.

To extend the model applicability to arbitrarily large images, Single GAN (SinGAN)^12^ and Single Denoising Diffusion Model (SinDDM)^13^ were proposed to learn the internal statistics of a given training image. Their shared coarse-to-fine architectural design enables the generation of image samples of any desired dimensions. Differing from single-image training, InfinityGAN ^14^ re-introduced large-scale training on patch-wise image data using low computational resources. Tailored for high-resolution natural scene creation, strong coordinate priors, such as vertical rapid saturation and horizontal repetitive patterns of sky, land, or ocean, were imposed within the structure and texture synthesizer of InfinityGAN.

In bioimage modeling, the utility of coordinate priors is nonetheless undesirable. This is because the arrangement of biological structures is not dictated by a rigid coordinate system, but rather by the intricate interplay between genetic, epigenetic, and gene expression variability that leads to the phenotype of a living system^15^. Taking spatial gene expression data as the input, we are motivated to propose **I**nfinite **S**patial **T**ranscriptomic **e**diting (IST-editing) that enables algorithmic gene expression-guided editing in a whole mouse pup. Importantly, the model training and inference can be efficiently executed on a single consumer-grade GPU, *e.g*., GeForce RTX 3090 Ti.

### Results

We test IST-editing on the public Xenium^16^ ST dataset of a one-day mouse pup (See also ‘Data availability’). This gigapixel-resolution dataset provides a well-curated sparse 3D array of 379-plex gene transcript counts (App. Fig. 3) and the spatially matched DAPI-stained WSI at identical resolution, offering a comprehensive morpho-molecular landscape of the living system.

#### Evaluation of generation results

We benchmark IST-editing (See also ‘Methods’) against state-of-the-art diffusion- and GAN-based models such as InfinityGAN. Consistent with the IST-editing approach, which utilizes densely and randomly sampled ST data pairs for training (Figure 1 (a)), we feed all the models with patch-wise spatial gene expression data (input) and DAPI images (output) for systematic and fair comparisons. Following the single-image training paradigm, we train SinGAN^12^ and SinDDM^13^ on individual tissue-level images (*e.g*., 4096×4096) and generate high-resolution images for direct comparison with the IST-editing results. In contrast, StyleGAN2, InfinityGAN, and IST-editing are trained on patch-wise data pairs extracted from the entire WSI. As shown in Figure 1 (b), SinGAN and SinDDM can recreate low-resolution images (small inset, left) including texture similarities to the original tissue such as the alveolar pattern observed in samples from the lung region. However, the image generation cannot be consistently scaled to a higher resolution (right): Only basic and biologically meaningless tissue textures remain. StyleGAN2 preserved a pattern resembling cell nuclei in generated high-resolution images. However, the tissue structure corresponding to the individual organ regions is lost, as is evident from the ‘StyleGAN2’ column of Fig. 1 (b). Owing to undesired coordinate priors for bioimage generation, we observed horizontal lines and repetitive patterns in images generated by InfinityGAN and clearly identifiable tissue structures are not present in these image examples.

**Figure 1.**
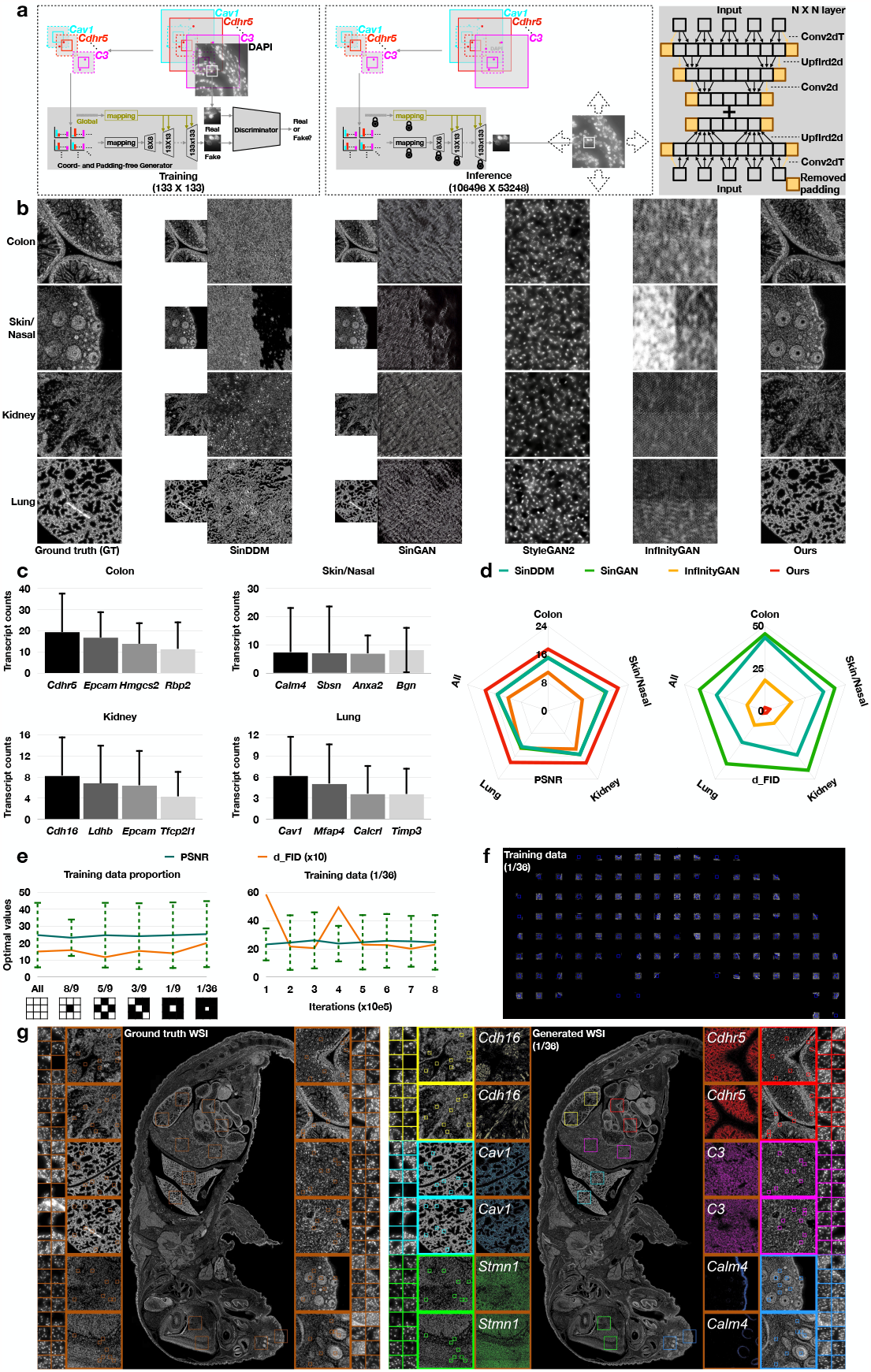
Model architecture and WSI generation. **a**. The conceptual illustrations of the proposed model. **b**. The visual comparison of tissue-level (4096×4096) synthesized images obtained by training with 100% of the available data using SinGAN^12^, SinDDM^13^, StyleGAN2^9^ and InfinityGAN^14^ as compared to IST-editing. **c**. The mean and standard deviation of transcript counts of the highly expressed genes (per cell) w.r.t. individual tissue regions. **d**. The comparison of tissue-level generation results between the compared methods by Peak Signal-to-Noise Ratio (PSNR) (left) and Fréchet Inception Distance *d*_FID_ (right). **e**. The comparison of PSNR and *d*_FID_ scores obtained by training IST-editing on progressively smaller subsets of available data (left) and at different numbers of iterations for training with 3% of the available data (right). For these experiments, subsets of the available data are sampled following the ‘checkerboard’ patterns, as illustrated underneath the ‘Training data proportion’ plot. **f**. The visual illustration of 3% of available training data. **g**. The cell-, tissue- and animal-level visualization of ground-truth (left) and generated (right) mouse pup WSI. To visualize the spatial pattern of leading gene expressions in the right plot, we first downscale the resolution of gene expression array using sum reduction and then shift the gene expression level to [0, 255].

After inputting the 379-plex gene expression data *, our approach successfully generates tissue-level images at the scale of 4096×4096 resolution, with biologically meaningful details (Fig. 1 (b), right). The generated images show a high level of similarity both in tissue organization, texture and cell-level detail to the biological prior, as supported by expert pathologist interpretation. Quantitatively illustrated in Fig. 1 (d), IST-editing outperforms compared methods in terms of low Fréchet Inception Distance (*d*_FID_)^18^ and high Peak Signal-to-Noise Ratio (PSNR) score. Using the padding-free convolutional operations and matched boundaries of gene expression arrays between neighboring tiles, IST-editing further achieved the WSI generation with a 106496×53248 pixel resolution. Please see also App. Fig. 1 for more elaborated visualization.

#### Training data utility (100% - 3%)

Next, we evaluate the generation robustness of the proposed approach under conditions of increasing data scarcity. For this purpose, we utilize progressively smaller subsets of the available data for training. As depicted in Fig. 1 (e, left), the optimal *d*_FID_ and PSNR scores remain consistent as the amount of available data decreases. Only when reducing the training data to 1*/*36 of the original size (Fig. 1 (f)) do we start to observe a mild degradation in quantitative performance by *d*_FID_. Upon comparing the cell-, tissue- and animal-level generation quality achieved by training on the entire dataset (App. Fig. 1 (a, b)) and 3% (Fig. 1 (g, right)) of the available data, the visual discrepancy between the two gigapixel-resolution WSIs appears marginal, substantiating the adaptability of IST-editing to limited data scenarios, requiring the seamless synthesis of more than 97% of the unseen data.

#### Evaluation of editing effects

We investigate gene expression-guided IST-editing of WSI data by three distinct strategies. Experiments are performed on the generated in-silico mouse model which contains co-profiled ST and WSI data of all major mammalian organ systems.

1. **Direct scaling of gene expressions**. Organized structures of diverse tissue regions emerge when progressively scaling the expression levels of the top four genes by a factor of 0.5, 1 (baseline), and 2 (Fig. 2 (b, left and middle)), while remaining gene expressions are zeroed out. Such targeted editing is driven by the observable dominant impact on the morphological generation of the top four leading expressed genes. Interestingly, the editing effects exhibited biologically explainable heterogeneity across the different regions. In the colon section, we observe the emergence of internal epithelial structures orchestrated by the upscaling of leading genes including Epithelial Cell Adhesion Molecule, *EPCAM*. As muscle-specific genes are not represented in the top colon gene sets, the outer muscle layer remains absent in the reconstruction. In other examples, the clear structures and organizations of the lung region have been recovered by our approach, closely resembling the GT lung image, and image artifacts (*e.g*.,, white fluff on GT WSI scan) are effectively eliminated in the reconstruction, as these are not captured in the GT ST data. Calculated on the proportional ratio between edited and GT tissue regions highlighted in the bounding boxes (*e.g*., Fig. 2 (d)), the radar charts in Fig. 2 (b, right) demonstrate a consistent increase in cell-level metrics approaching the GT with the up-scaling coefficients.
2. **Indirect scaling of gene expressions**. Similar to the cell-level manipulation study^8^, we perform algorithmic editing on the sample covariance matrix (SCM) and scale the leading eigenvalues by 0.1, 1 (baseline), and 2. Consider the SCM 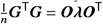, where ***G*** is the collection of *n* 379-plex gene expression data from a given tissue region, ***O***_*i*_ is the 379×379 eigenbasis and ***λ*** is the (sorted) diagonal eigenvalues derived from eigenvalue decomposition. Then, we control ***λ*** for indirectly conducting gene expression-guided editing. As illustrated in Fig. 2 (c), there exists a rather homogeneous transition of tissue structures across the various regions of interest. On the contrary to the results described above using the leading genes, the muscle layer of the colon tissue as well as global architectural features of lung and skin are already observed at the scale of 0.1 when using all genes as an input. When examining the editing effects with up-scaling of the eigenvalues, we witness a further increase in DAPI pixel intensity and increased sharpening of architectural details closely resembling the GT image. This is reflected by the quantitative analysis of the interpretable morphological features Fig. 2 (c, right), where we observe an expected increase in the cellular region and DAPI signals.
3. **Interpolation between unorganized and well-organized gene expressions**. To simulate morphological transitions at scale of the whole in-silico mouse model, we conduct linear interpolation between randomly sampled and ground truth spatial gene expressions, generating WSI results at coefficients of 0 (noise), 0.5, and 1 (mouse pup). The resulting WSIs exhibit a gradual progression from chaotic cellular organization - as reflected through the appearance of ‘random noise’ across the entire sample - to the highly organized structure of the one-day mouse pup. Together with the above editing experiments, we demonstrate the versatility of IST-editing in modeling biological processes across multiple biological scales.

#### Evaluation of failure cases and discussion

Pushing the limits further, we conduct extreme stress tests on the proposed approach for reconstructing the whole organism. This is carried out by training on a single 2048×2048 resolution image extracted from individual tissue regions such as kidney, lung, and brain. Though the overall outline and structure of the mouse pup are retained, IST-editing struggles to recreate the WSI with fine biological-aware details, as illustrated in App. Fig. 4. Remarkably, heterogeneous generation patterns for different organs arise when training solely on one single image. For instance, the training of the gut region image leads to the ‘black hole’ generation of the mouse brain. This can be explained by the non-overlapping highly expressed genes between the gut (*e.g*., *Cdh16, Ldhb, Epcam, Tfcp2l1*) and brain (*e.g*., *Stmn1, Gap43, Nnat, Tubb3*) region, as presented in Fig. 2 (a) and App. Fig. 3. A similar pattern occurs when utilizing the liver tissue image with matched high-expressed genes such as *Cav1, Mfap4, Calcrl* and *Timp3*. When utilizing an ‘almost black’ image with a mere fragment of mouse skin (‘Skin’ of App. Fig. 4), the overall structure of the mouse pup remains preserved, though the cellular and tissue generation tends to exhibit a preference for mimicking skin epithelial morphology, suggesting a bias towards replicating trained cellular subtypes.

**Figure 2.**
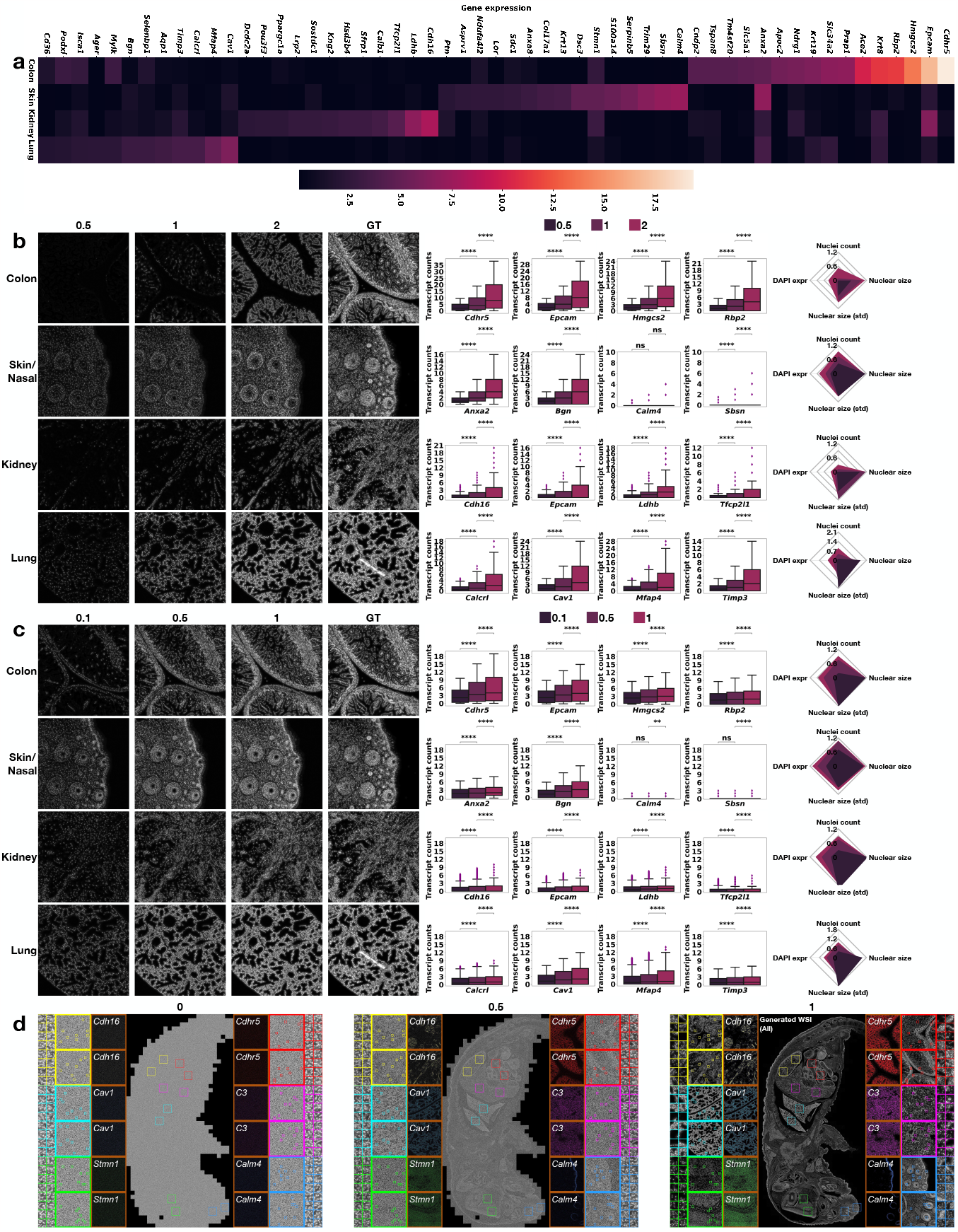
Gene expression profiles and experimental results of IST-editing. **a**. The heatmap of highly expressed genes (average per cell) w.r.t. different tissue regions of the whole mouse pup and selected organ systems of interest. **b**. The visual (left) and quantitative (right) IST-editing effects on individual tissue regions obtained by scaling the leading gene expressions (middle) while zeroing out the rest of gene expressions. **c**. The visual (left) and quantitative (right) IST-editing effects on individual tissue regions obtained by scaling the leading eigenvalues of the sample covariance matrix (SCM)^8,17^. For both (b) and (c), **** means *p* ≤ 0.0001 and ‘ns’ stands for not statistically significant. The error bar of the box plot represents the 5%–95% quantile. The radar plots report the proportional ratio of morphological features between edited (numerator) and GT cells (denominator). **d**. The overall editing effects on the whole mouse pup achieved by the interpolation between random noise and ground-truth gene expressions. For plots (b)-(d), all the editing experiments are conducted using the model trained with 100% of the available data.

Utilizing the DAPI-stained WSI that captures nucleus morphology, this proof-of-concept study showcased the generative ability and editability in a generated living system. Incorporating additional staining techniques that strongly increase the detail of cellular components, such as cytoplasm, cell membranes, and extracellular matrix, IST-editing can be employed to further untangle complex phenotypic traits of living organisms. This is, for instance, supported by the first Hematoxylin and Eosin (H&E) generation results of the same mouse pup (Fig. App. 5). The potential of IST-editing extends beyond animal modeling. By adapting the proposed approach to the analysis of samples from human pathology, IST-editing can provide a novel perspective to investigate the linkage between genotype and phenotype in human diseases.

## Methods

To efficiently process the paired transcript count array and DAPI image with matched gigapixel resolutions, we develop IST-editing upon the StyleGAN2 backbone. This is motivated by recent publications on GAN studies^11,19,20^ in response to the remarkable advances made by diffusion models. While being orders of magnitude faster at inference time, these methods, built upon advanced GAN architectures such as StyleGAN2, exhibit superior generation and editing performance that remain competitive with their diffusion counterparts. Notably, IST-editing can be effortlessly adapted to various spatial multi-omics technologies that enable concurrent profiling of spatially resolved molecular and bioimage data.

### Training data

In natural image generation, previous generative models^12,13^ typically utilize (spatial) noise input for unconditional image generation. In addition, learned textural representations^21^ can be incorporated into the model to guide the image alterations^2^. However, semantic ambiguity often occurs in interpreting a single latent code and qualitative analysis is mostly made possible for a subset of representations^22^. Given the well-established biological understanding of many individual genes, we take gene expression data as (latent) representation(s) and input to guide bioimage generation. To alleviate the data loading bottleneck for processing an entire WSI, we divide the 3D gene expression array and WSI into smaller tiles with 2176×2176 pixel resolution, which share overlapped 2176×64 boundary regions with neighboring tiles. As shown in Fig. 1 (a), we then utilize the patch-wise spatial gene expression and DAPI image for training, where the training pairs are randomly and densely sampled from the processed tile data. Specifically, the cropped 3D gene expression array has 256×256×379 resolution, which is center-aligned to the paired 128×128 DAPI image. A higher resolution of the gene expression array is crucial for preserving the boundary consistency between generated neighboring tiles. Due to the high spatial sparsity, we down-scale the gene expression array to 8×8×379 by sum reduction along spatial dimensions, such that more densely distributed gene expressions are aggregated in the format of a smaller 3D array.

### Coordinate- and padding-free generator *G*

Instead of relying on strong coordinate-based priors, such as the vertical saturation and horizontal repetition of natural scenes, the design of our generator is driven by the intricate interaction between genes (causative factors) and phenotypes (observable characteristics). To model the directed linkage from gene expression to the DAPI image, we propose a straightforward coordinate-free generator, which is constructed using a series of padding-free StyledConv layers. No external prior knowledge, aside from gene expression data, is incorporated into the output DAPI images. In all padding-free layers, we discard pixel values that are padded at both spatial ends of the output. As a result, we have consequential image outputs with 8, 13, 21, 37, 69, and 133 spatial resolutions. After discarding 5 boundary pixels of the last-layer output, we output the 128×128 DAPI image. It is worth mentioning that, by appending more padding-free layers, our model can be easily extended to the generation of higher resolution images such as 256 × 256 and 512 × 512.

### Cell-subtype conditioned discriminator *D*

Inspired by conditional generations of well-characterized normal and cancer cellular images^8^, we aim to integrate cell subtype information into the discriminator to adversarially and conditionally train the generator. In the absence of clear cell-level annotations in the Xenium dataset, we conducted a careful evaluation of the WSI and cell-level clustering* within the context of tissue organizations. Instructed by domain biomedical experts, we confirmed the accuracy of subtype assignments derived from the ‘kmeans_10_clusters’ results in the raw data. Depending on the majority vote of cell subtypes presented in the sampled training data, we assign the label of the predominant subtype to each image tile. Thereafter, we integrate cell label embeddings into *D* and train both models with the conditional adversarial loss ℒ_adc_. Along with the *R*_1_ regulation ℒ_*R*1_ and path length regulation loss ℒ_path_^9^, we have the following loss function

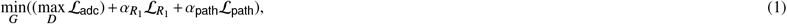

where 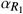 and *α*_path_ are hyperparameters. Then, we train the GAN model for 800k iterations with a batch size of 16. Eventually, the optimal model performance is determined using *d*_FID_ and PSNR. For the former, we use a more efficient implementation^17^ and robust CLIP features^21,23^ to carry out the computation.

### Inference

At inference, we divide the WSI generation into patch-wise image generations that are parallelizable on the GPU. In particular, the boundary consistency of neighboring generated tiles is guaranteed by the gene expression arrays with overlapped 256×64 boundary regions and seamless window sliding of padding-free StyledConv layers. Using a single GeForce RTX 3090 Ti, it takes ∼ 30 mins to synthesize a 106496×53248 WSI. The generated DAPI and H&E WSIs are accessible via the repository link and can be thoroughly examined using open-source software such as QuPath^24^.

## Data availability

The Xenium ST dataset includes the spatially resolved 379-plex mRNA transcripts and the matched gigapixel DAPI-stained WSI, in which 1.36 million cells are identified and segmented for downstream analysis. The dataset is accessible via https://s3-us-west-2.amazonaws.com/10x.files/samples/xenium/1.6.0/Xenium_V1_mouse_pup/Xenium_V1_mouse_pup_outs.zip, The downloading links to the dataset and our generated WSIs are free and publicly available to users, without additional login or registration requirements.

## Author contributions statement

J.W. and V.H.K. conceived the research idea. J.W. implemented the algorithm and carried out the experiments. J.W., I.B. and V.H.K. analyzed the results. J.W. and V.H.K. drafted the manuscript. I.B. critically reviewed the manuscript and supplied biological interpretations. V.H.K. supervised the project.

## Acknowledgements

This study is funded by core funding of the University of Zurich to the Computational and Translational Pathology Lab led by V.H.K. at the Department of Pathology and Molecular Pathology, University Hospital and University of Zurich.

## Footnotes

* Please see also see the 10x Genomics data summary provided at https://cf.10xgenomics.com/samples/xenium/1.6.0/Xenium_V1_mouse_pup/Xenium_V1_mouse_pup_analysis_summary.html.

## References

1. Moses, L. & Pachter, L. Museum of spatial transcriptomics. Nat. Methods 19, 534–546 (2022).

2. Bermano, A. H. et al. State-of-the-art in the architecture, methods and applications of stylegan. In Computer Graphics Forum, vol. 41, 591–611 (Wiley Online Library, 2022).

3. Croitoru, F.-A., Hondru, V., Ionescu, R. T. & Shah, M. Diffusion models in vision: A survey. IEEE Transactions on Pattern Analysis Mach. Intell. (2023).

4. Royer, L. A. The future of bioimage analysis: a dialog between mind and machine. Nat. Methods 20, 951–952 (2023).

5. Carrillo-Perez, F. et al. Synthetic whole-slide image tile generation with gene expression profile-infused deep generative models. Cell Reports Methods 3 (2023).

6. Lamiable, A. et al. Revealing invisible cell phenotypes with conditional generative modeling. Nat. Commun. 14, 6386 (2023).

7. Wu, J. & Koelzer, V. H. Gilea: Gan inversion-enabled latent eigenvalue analysis for phenome profiling and editing. bioRxiv 2023–02 (2023).

8. Wu, J. & Koelzer, V. H. In silico spatial transcriptomic editing at single-cell resolution. bioRxiv (2023).

9. Karras, T. et al. Analyzing and improving the image quality of stylegan. In Proceedings of the IEEE/CVF conference on computer vision and pattern recognition, 8110–8119 (2020).

10. Karras, T. et al. Alias-free generative adversarial networks. Adv. Neural Inf. Process. Syst. 34, 852–863 (2021).

11. Kang, M. et al. Scaling up gans for text-to-image synthesis. In Proceedings of the IEEE/CVF Conference on Computer Vision and Pattern Recognition (2023).

12. Shaham, T. R., Dekel, T. & Michaeli, T. Singan: Learning a generative model from a single natural image. In Proceedings of the IEEE/CVF international conference on computer vision, 4570–4580 (2019).

13. Kulikov, V., Yadin, S., Kleiner, M. & Michaeli, T. Sinddm: A single image denoising diffusion model. In International Conference on Machine Learning, 17920–17930 (PMLR, 2023).

14. Lin, C. H., Lee, H.-Y., Cheng, Y.-C., Tulyakov, S. & Yang, M.-H. Infinitygan: Towards infinite-pixel image synthesis. In International Conference on Learning Representations (2022).

15. Haniffa, M. et al. A roadmap for the human developmental cell atlas. Nature 597, 196–205 (2021).

16. Janesick, A. et al. High resolution mapping of the breast cancer tumor microenvironment using integrated single cell, spatial and in situ analysis of ffpe tissue. bioRxiv 2022–10 (2022).

17. Wu, J. & Koelzer, V. Sorted eigenvalue comparison dEig: A simple alternative to dFID. In NeurIPS 2022 Workshop on Distribution Shifts: Connecting Methods and Applications (2022).

18. Heusel, M., Ramsauer, H., Unterthiner, T., Nessler, B. & Hochreiter, S. Gans trained by a two time-scale update rule converge to a local nash equilibrium. Adv. neural information processing systems 30 (2017).

19. Sauer, A., Schwarz, K. & Geiger, A. Stylegan-xl: Scaling stylegan to large diverse datasets. In ACM SIGGRAPH 2022 conference proceedings, 1–10 (2022).

20. Sauer, A., Karras, T., Laine, S., Geiger, A. & Aila, T. Stylegan-t: Unlocking the power of gans for fast large-scale text-to-image synthesis. Int. Conf. on Mach. Learn. (2023).

21. Radford, A. et al. Learning transferable visual models from natural language supervision. In International conference on machine learning, 8748–8763 (PMLR, 2021).

22. Härkönen, E., Hertzmann, A., Lehtinen, J. & Paris, S. Ganspace: Discovering interpretable gan controls. Adv. neural information processing systems 33, 9841–9850 (2020).

23. Kynkäänniemi, T., Karras, T., Aittala, M., Aila, T. & Lehtinen, J. The role of imagenet classes in fr\’echet inception distance. arXiv preprint arXiv:2203.06026 (2022).

24. Bankhead, P. et al. Qupath: Open source software for digital pathology image analysis. Sci. reports 7, 1–7 (2017).

