## Supplemental Figure 1 - 5 for "IST-editing: Infinite spatial transcriptomic editing in a generated gigapixel mouse pup"

### ABSTRACT

Advanced spatial transcriptomics (ST) techniques provide comprehensive insights into complex living systems across multiple scales, while simultaneously posing challenges in bioimage analysis. The spatial co-profiling of biological tissues by gigapixel whole slide images (WSI) and gene expression arrays motivates the development of innovative and efficient algorithmic approaches. Using Generative Adversarial Nets (GAN), we introduce **I**nfinite **S**patial **T**ranscriptomic **e**diting (IST-editing) and establish gene expression-guided editing in a generated gigapixel mouse pup. Trained with patch-wise high-plex gene expression (input) and matched image data (output), IST-editing enables the synthesis of arbitrarily large bioimages at inference, *e.g.*, with a $106496 \times 53248$ resolution. After feeding edited gene expressions to the trained network, we model cell-, tissue- and animal-level morphological transitions in the generated mouse pup. Lastly, we discuss and evaluate editing effects on interpretable morphological features. The generated WSIs of the mouse pup and code are publicly released and accessible via https://github.com/CTPLab/IST-editing.

### Appendix

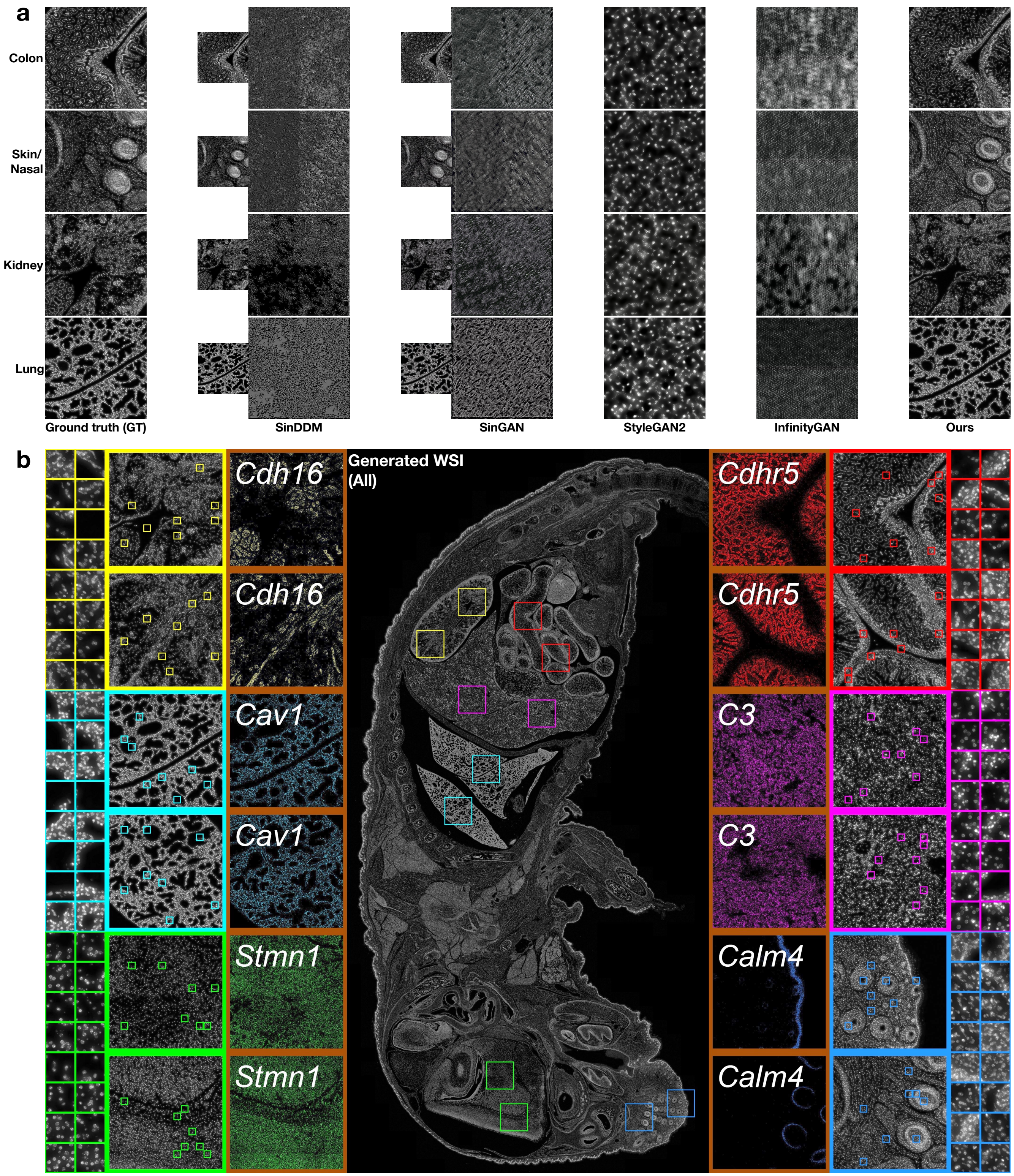


**Figure 1.** The generation results of tissue region images (a) and WSI (b).

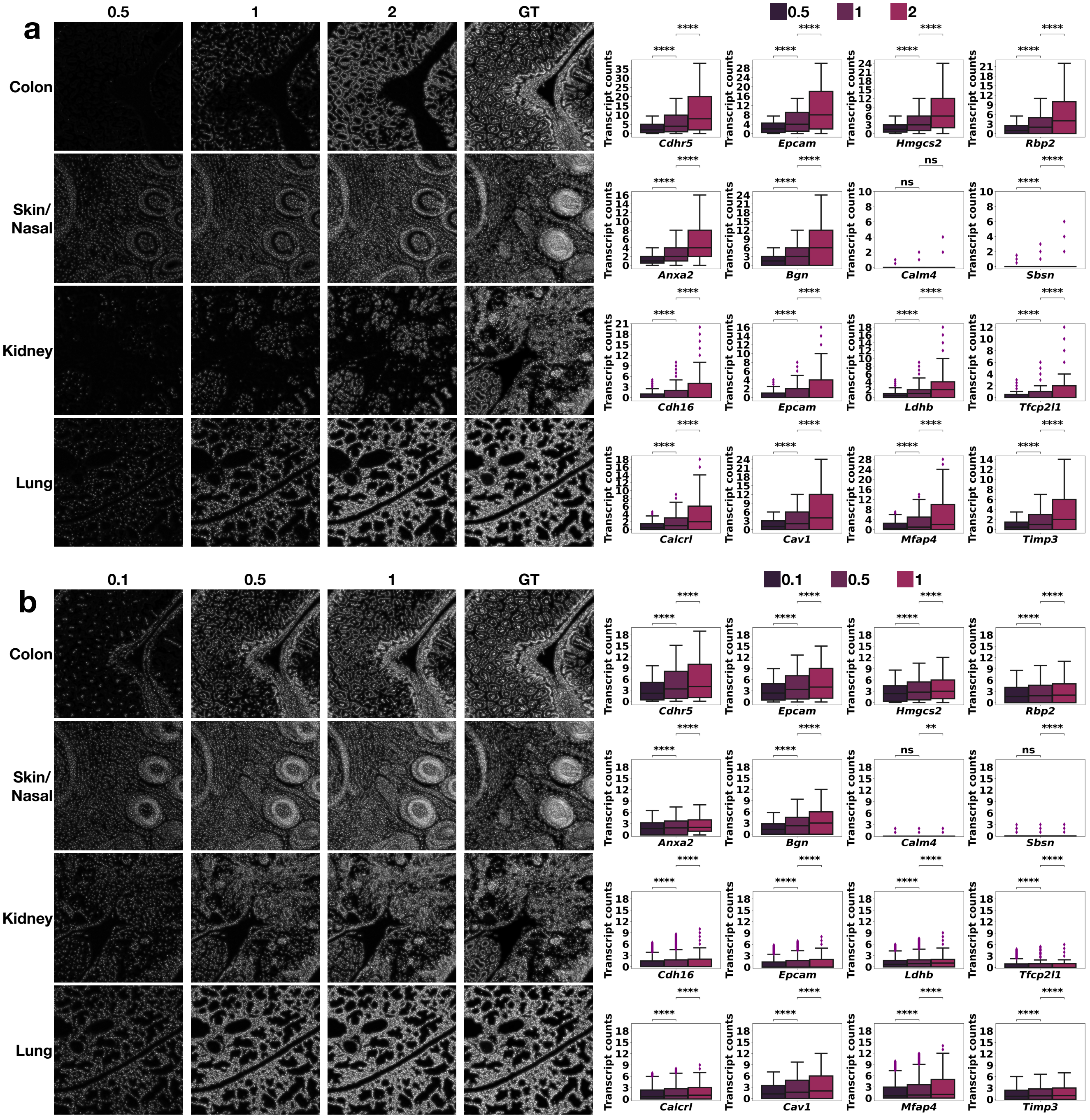


**Figure 2. The experimental results of diverse editing effects**. **a**. The visual (left) and quantitative (right) editing effects on various tissue regions by scaling the leading gene expressions (middle) while zeroing out the rest of gene expressions. **b**. The visual (left) and quantitative (right) editing effects by scaling the leading eigenvalues of the sample covariance matrix (SCM) of individual tissue regions.

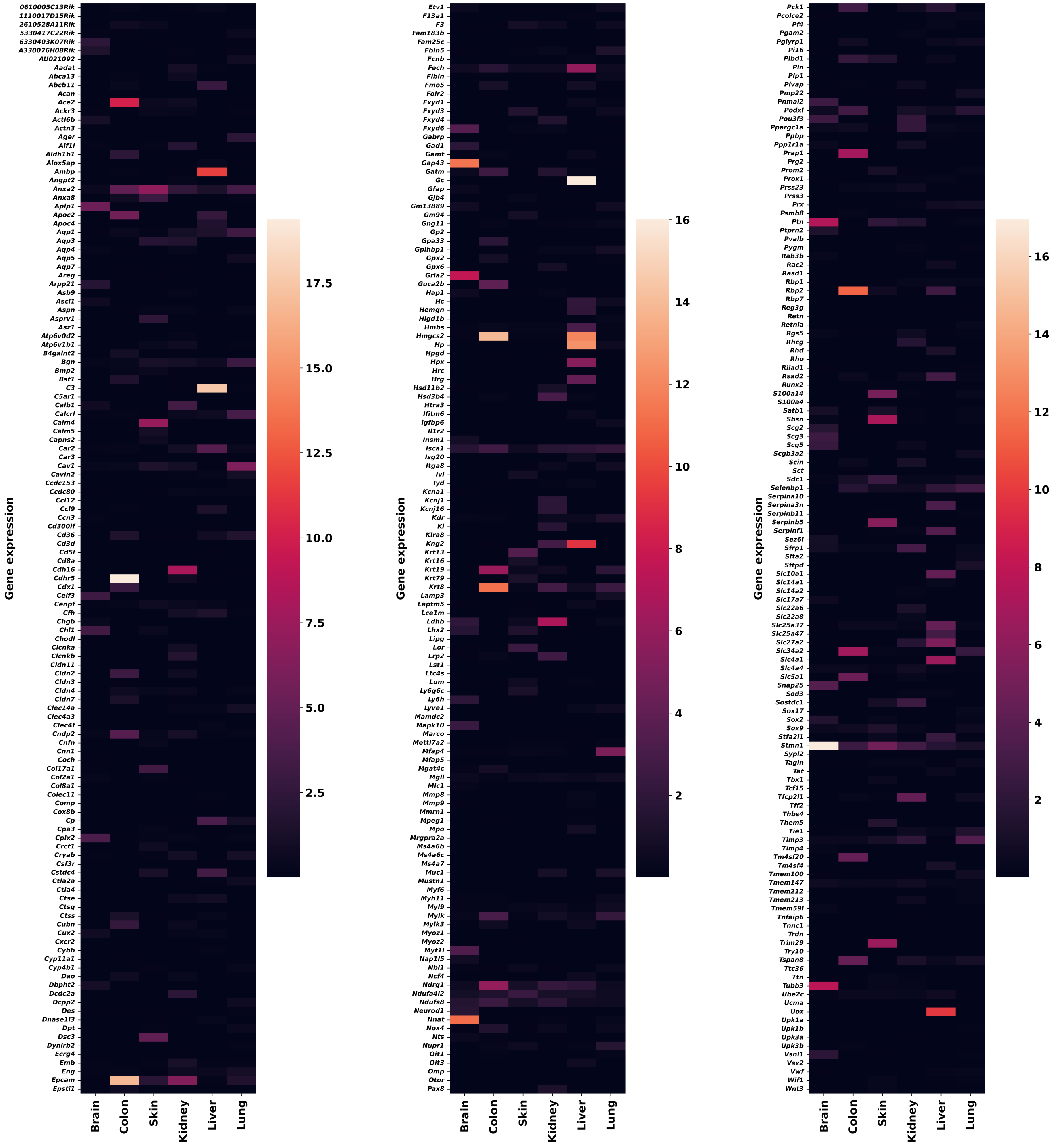

Figure 3. The heatmap of 379-plex gene expressions (average per cell) w.r.t. different tissue regions.

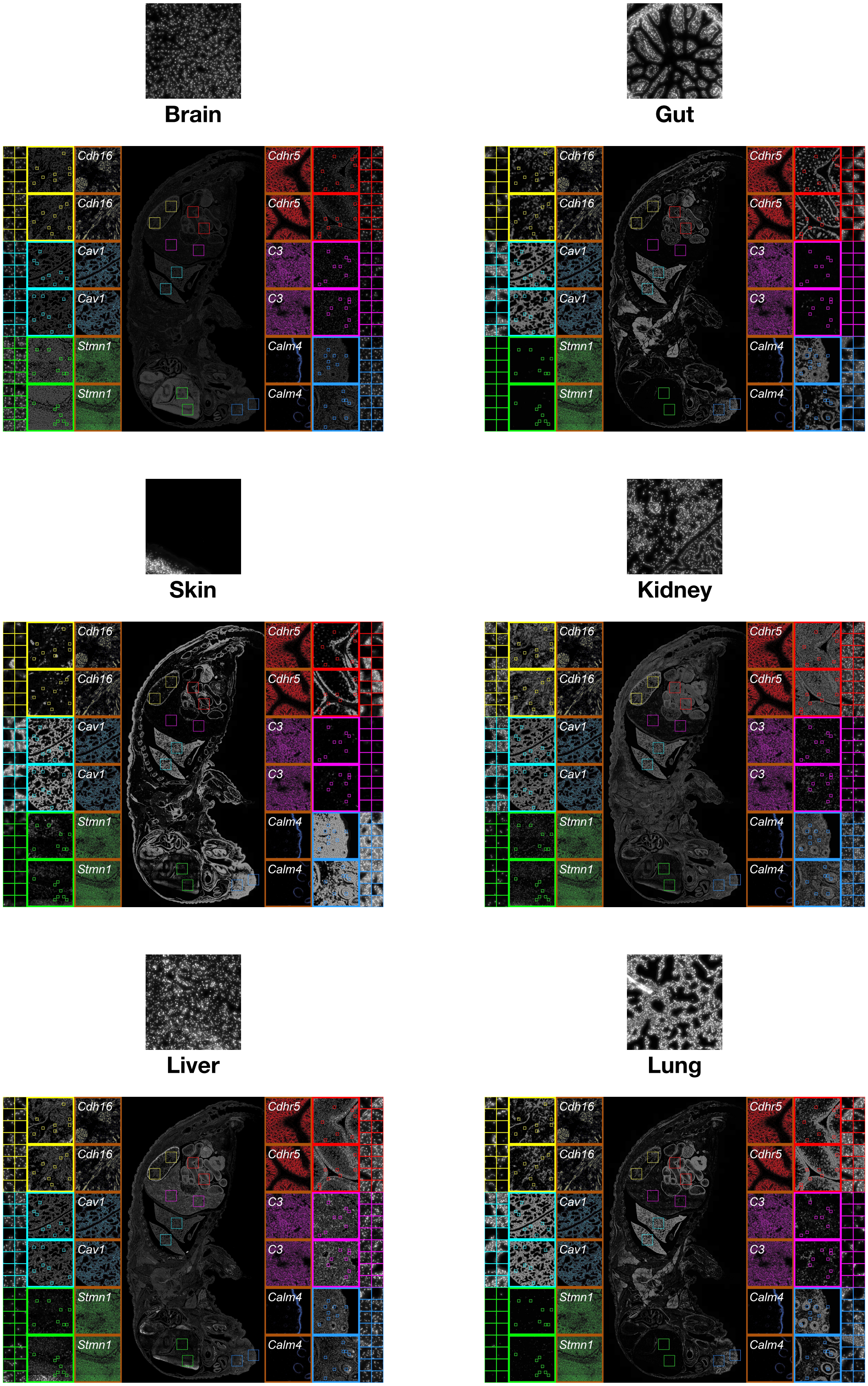


**Figure 4.** The failure cases of WSI generation brought by training on a single 2048 × 2048 image extracted from individual tissue regions.

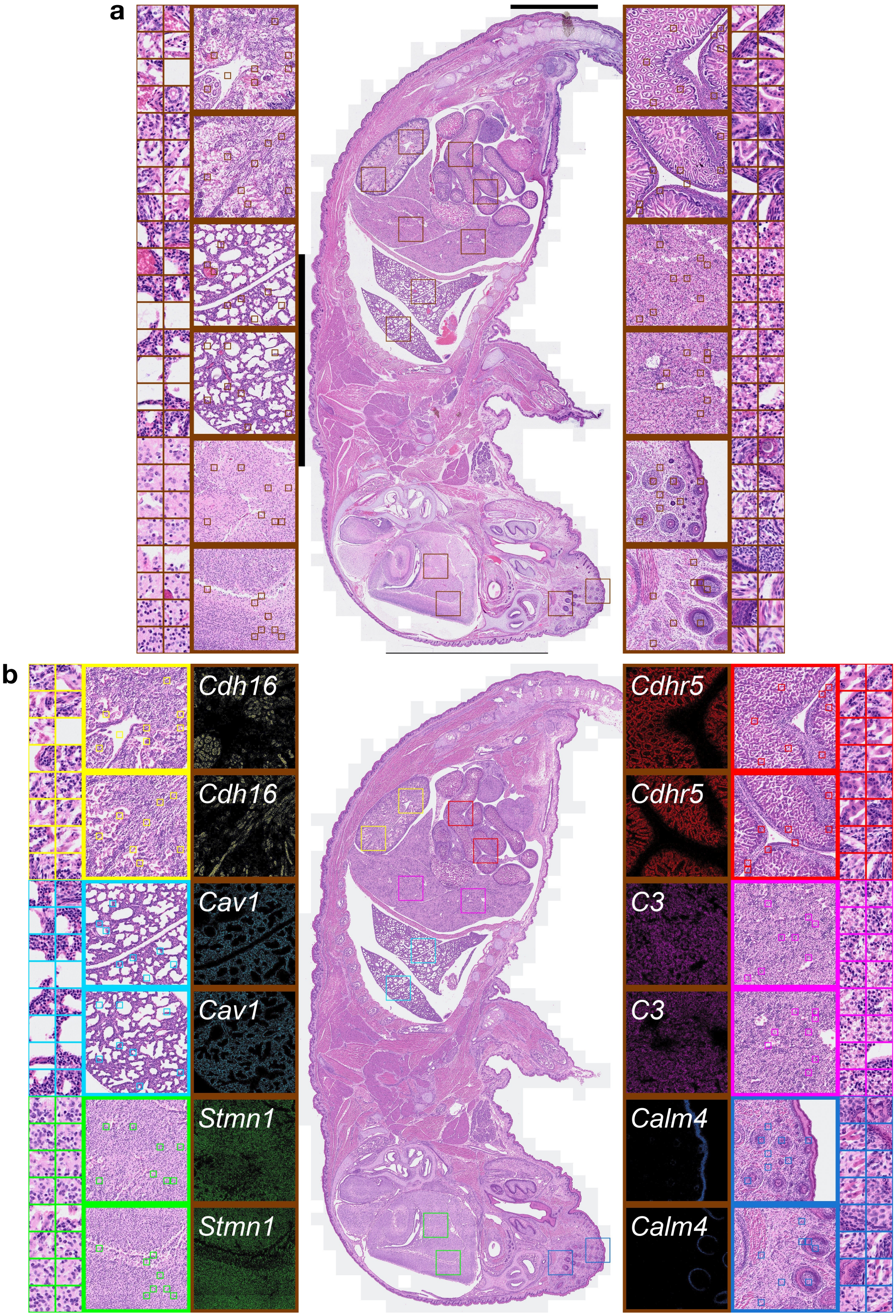


**Figure 5.** The ground truth (a) and generated (b) H&E-stained WSIs.
